# Assessment of the gene expression profiles of leukocytes from different isolations

**DOI:** 10.1101/678904

**Authors:** Yanru Xing, Xi Yang, Haixiao Chen, Sujun Zhu, Jinjin Xu, Yuan Chen, Juan Zeng, Fang Chen, Hui Jiang, Wen-Jing Wang

**Author notes:** Corresponding author: Wen-Jing Wang, address: Build 11, Beishan Industrial Zone, Yantian District, Shenzhen, 518083, China.; Hui Jiang, address: Build 11, Beishan Industrial Zone, Yantian District, Shenzhen, 518083, China. These authors contributed equally (Yanru Xing, Xi Yang).

## Abstract

**BACKGROUND:** Research on peripheral leukocyte gene expression in human health and disease is growing rapidly. However, how to process sample efficiently, simply and stably, and how to reflect human physiological state preferably remains critical issues in large cohort studies.

**METHODS:** We used RNA-seq to explore the differences of gene expression profiles among whole blood (WB) and three groups of leukocytes from buffy coat (BC) extraction, red blood cell (RBC) lysis and peripheral blood mononuclear cell (PBMC) isolation.

**RESULTS:** The residual globin mRNA in leukocytes from RBC lysis (1.00% ±1.23%) and PBMC isolation (0.06% ± 0.03%) was much less than that in leukocytes from BC extraction (17.48% ± 6.95%) and WB (24.46% ± 6.43%), resulting in higher transcriptome mapping rates and larger numbers of detected genes. The expression of 616 genes associated with leukocyte function was slightly higher in leukocytes from RBC lysis than that from BC extraction and WB, but barely detected in leukocytes from PBMC isolation.

**CONCLUSIONS:** We suggest that sample processing based on RBC lysis could allow better applications of gene expression profiling of peripheral leukocytes in large cohort studies.

## Background

Leukocytes in peripheral blood are a part of the immune system that helps fight infection and defend the body against other foreign materials. Leukocytes in different diseases including rheumatoid arthritis [1], type 2 diabetes [2], and infectious diseases [3] have specific characteristics of expression. Different environmental stresses, such as sports [4], and smoking [5] will also reflect in gene expression of Leukocytes. As peripheral blood is more available and less risky to collect than invasive organ biopsy, leukocyte is an ideal type of cells as biomarkers for the diagnosis. Besides that, as trans-omics approaches are applied to many cohort studies, leukocyte transcription plays an important role in it.

Whole blood (WB) keeps all types of blood components including red blood cells, leukocytes and platelets. While, globin mRNA ratio varies with red blood cells which make up around 45% of WB volume, and influence the RNA-seq result. Beyond that, differential expression analysis has showed that in the case of large percent of globin mRNA during RNA-seq, thousands of genes are significantly down-regulated, and the ability to detect real biological mutations has been reduced [6]. So, it is necessary to isolate leukocyte from peripheral blood. There are several methods to isolate leukocytes before RNA extraction. The buffy coat (BC) extraction mainly includes leukocytes and platelets, but, inevitably, there is some red blood cells contamination. The red blood cell (RBC) lysis removes red blood cells and leaves leukocytes and platelets [7]. Peripheral blood mononuclear cell (PBMC) isolation keeps all mononuclear leukocytes, such as lymphocytes and monocytes [8].

Expression profiles of leukocytes vary with isolations, while how to reflect human physiological state preferably in cohort studies is largely unknown. In this study, we used RNA-Seq to investigate gene expression profiles of leukocyte with different isolations. Our data suggests that differences caused by isolation should not be ignored before starting a large-scale cohort studies.

## Materials and Methods

### Sample collection

For the isolation assessment, fresh peripheral blood samples from 3 healthy volunteers (two males and one female) were collected using EDTA tubes and processed immediately. Three different isolations were used and WB as a control for each sample before RNA extraction (Figure 1A). The study protocol was approved by the BGI Institutional Review Board (NO. BGI-IRB 17034). All donors gave their written consent for non-therapeutic use of their donated blood samples.

**Figure 1:**
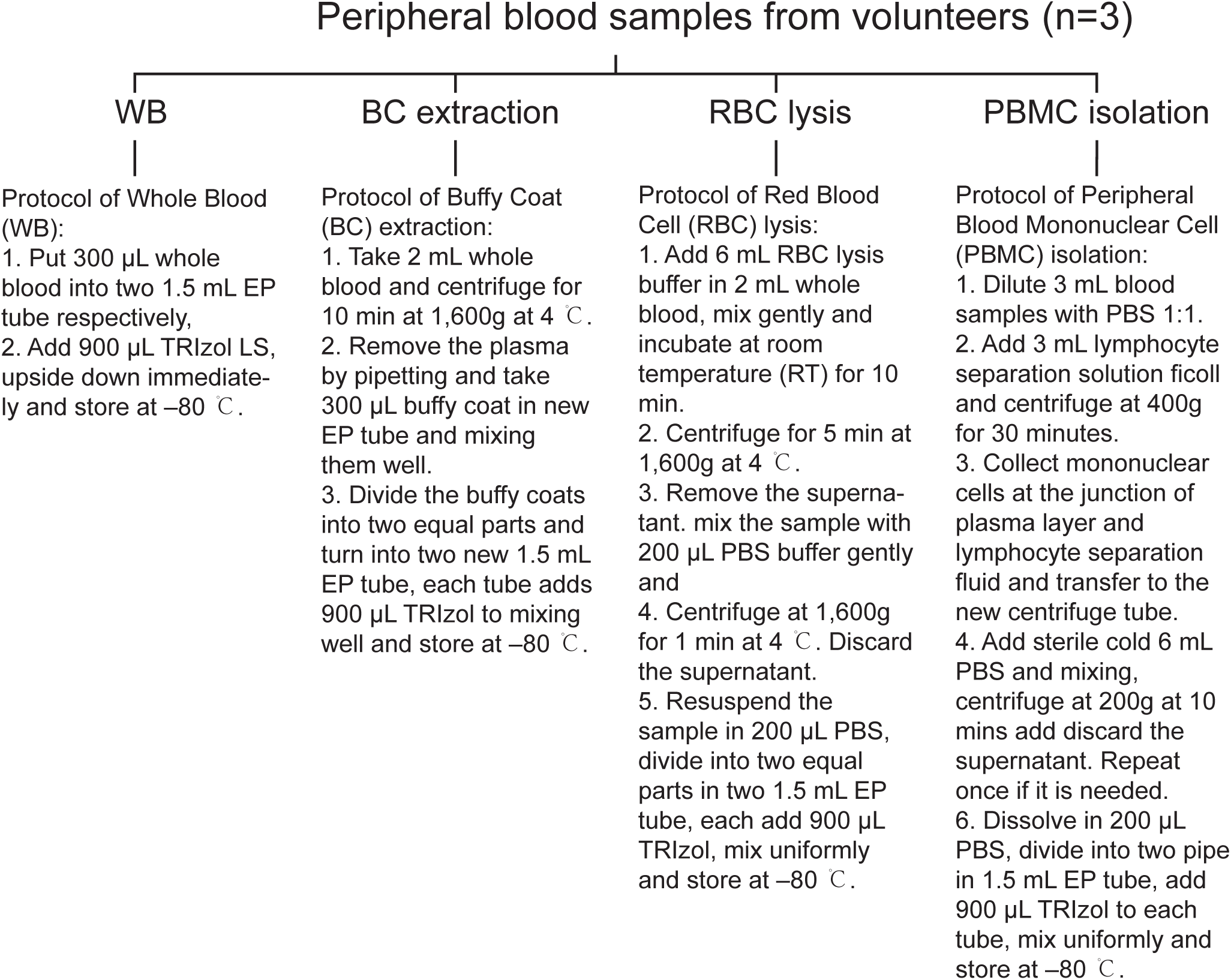
Study design. Protocol of WB and three leukocytes isolation methods.

### RNA extraction, library preparation and sequencing

Total RNA was extracted by the TRIzol Reagent (Thermo Fisher, 15596026) and TRIzol LS Reagent (Thermo Fisher, 10296028) according to manufacturer’s manual. RNA concentration and integrity were calculated by Agilent 2100 bioanalyzer (Agilent Technologies, G2939A) and accompanying software. Samples were considered to be acceptable if the total RNA was greater than or equal to 200ng and the RIN score was greater than or equal to 6. RNA Libraries were constructed through RNase H method [9]. Sequencing was performed on the BGISEQ-500RS (single end 50bp) developed by BGI.

### Data preprocessing

Soap2 [10] (Version 2.21t) was used to remove rRNA (hg19 rRNA ref), identify globin mRNA reads (globin list was shown in Supplementary Table S1) and calculate the proportion of rRNA, globin mRNA in raw reads. Then SOAPnuke [11] (Version 1.5.6) was used to filter reads and remove reads with adaptors. Reads with unknown bases (N) more than 10% and low-quality reads, which were defined as a read in which the percentage of the base is less than 5, were greater than 50%, were filtered. Clean data were obtained in FASTQ format. Finally, Bowtie2 [12] (Version 2.2.5) was used to align to the known gene transcriptome annotations from the UCSC Genome Browser [13] and generate bam files for analysis. Each read was mapped once to the best possible mapping location with default parameters. In the instance of a tie between two or more mapping locations, the read would be randomly assigned to one of the tied locations. The Bowtie2 output SAM file was converted into the BAM format using the SAMtools [14] option.

### Differentially expressed gene analysis

The gene expression abundance was quantified by RSEM [15] package at gene level. We kept genes with at least one count per million (CPM) in at least six samples. TMM normalization was performed to eliminate composition biases between libraries. We used edgeR [16] and QL F-test to identify DEGs for two-group comparisons: any two pretreatments of peripheral blood (WB, BC extraction, RBC lysis and PBMC isolation). The Benjamini-Hochberg (BH) method [17] was employed to estimate false discovery rate and correct multiple comparisons. DEGs were considered significantly associated with two groups if they exhibited a BH-adjusted P value<=0.01 and a log2 fold change>=1. MFUZZ [18] of R was used to implemented soft clustering for DEGs’ gene expression (Log2(RPKM+1) transformation was performed) in isolation test.

### Gene function analysis

Functional annotation of genes obtained from edgeR was performed by Metascape [19]. Genes information was searched in GeneCards [20] and Ensembl database [21].

### Statistics

For two-group and multigroup comparisons, we used unpaired two-sided t-test or multigroup analysis of variance (ANOVA), respectively. Group data are presented as mean ± sd. Lm function in R was used to fit linear models between globin mRNA and mapping ratio or detected gene number, figure showed level of confidence interval with 99%. To determine associations of two group gene expression, we used coefficient of determination.

## Results

### RNA quality and RNA-seq data information

All samples were extracted and the RNA library successfully constructed (Supplementary Table S2). For each sample of RNA-Seq, about 100M clean reads were obtained respectively (Supplementary Table S3).

### Assessment of globin mRNA depletion in different isolations

As preview studies showed that abundant globin mRNA influences RNA-seq analysis [22, 23], we assessed the three leukocytes isolations, and WB as a control. We found globin mRNA ratio was performed WB > BC extraction > RBC lysis ≥ PBMC isolation (Figure 2A), and significantly different among them (p= 4.96e-04, Supplementary Table S4). The BC extraction -- which isolated the leukocytes of peripheral blood only -- had a much higher globin mRNA ratio (17.48% ± 6.95%) than RBC lysis (1.00% ±1.23%) and PBMC isolation (0.06% ± 0.03%). Transcriptome mapping ratio and detected gene number were performed PBMC ≥ RBL > BC > WB (Figure 2B and Figure 2C). We also found globin mRNA ratio had high influence on proportion of transcriptome (p-value: 5.737e-04, Adjust R-squared: 0.68) and detected gene number (p-value: 9.746e-05, Adjust R-squared: 0.77) (Figure 2D, Figure 2E).

**Figure 2:**
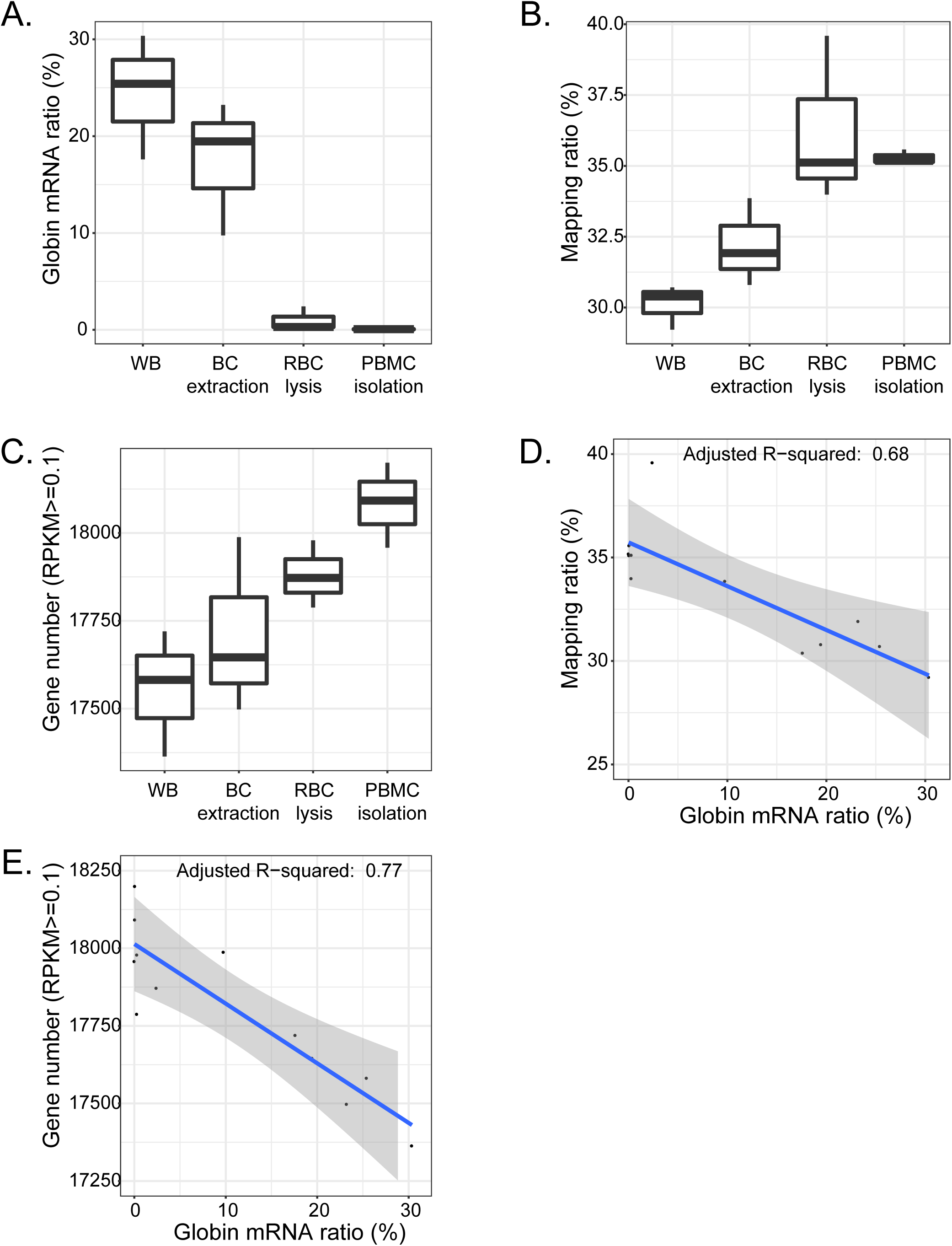
Globin mRNA ration and gene detected ability in WB and three leukocytes isolation methods. (A) Percentage of reads mapping to globin mRNA. WB (24.46% ± 6.43%, means ± standard deviation), BC extraction (17.48% ± 6.95%), RBC lysis (1.00% ±1.23%), PBMC isolation (0.06% ± 0.03%). (B) Proportion of reads mapping to transcripts. WB (30.11% ± 0.78%), BC extraction (32.19% ± 1.55%), RBC lysis (36.23% ± 2.96%), PBMC isolation (35.29% ± 0.26%). (C) Gene number (RPKM >= 0.1). WB (17555 ± 179), BC extraction (17711 ± 251), RBC lysis (17880 ± 96), PBMC isolation (18083 ± 121). (D) Relationship between the globin mRNA ratio and mapping ratio. (E) Relationship between the globin mRNA ratio and gene number. Gray shadow means a level of 99% for the confidence interval.

### Gene expression profiles influenced by different isolations

The difference in gene expression caused by isolations was greater than the difference between individuals (Figure 3A). However, one of BC extraction samples was clustered in RBC lysis group and the other two were clustered in WB group; we found that the sample clustered in RBC lysis group had a lower globin mRNA ratio (9.76%) than samples clustered in WB group (BC extraction_2 with 19.46%, BC extraction_3 with 23.23%). Although BC extraction get rid of parts of red blood cells in the process, gene expression profiles were similar between BC extraction and WB, and no DEG was found (Figure 3B, Figure 3C).

**Figure 3:**
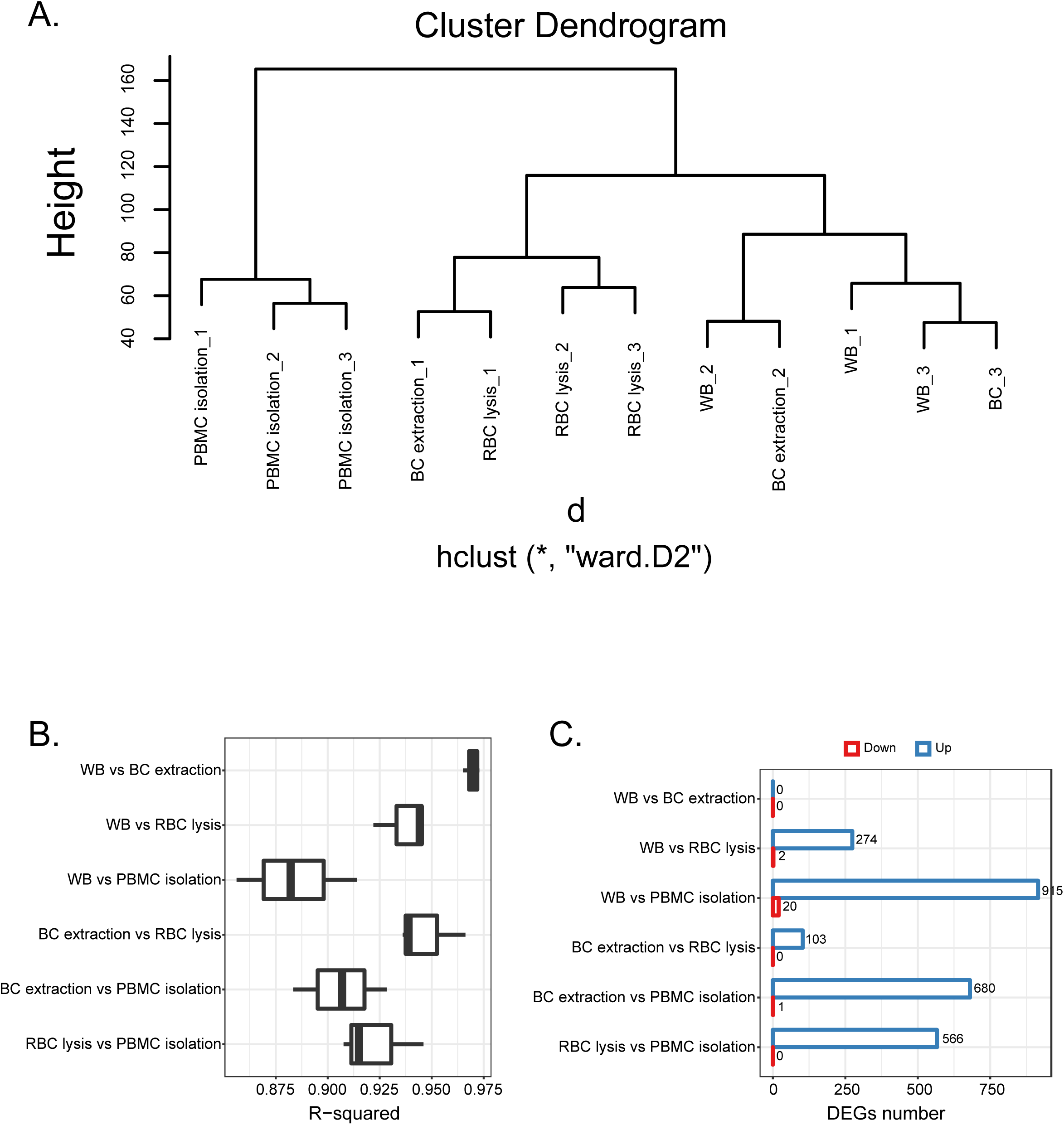
Consistency and difference of gene expression among different groups. (A) Hierarchical clustering dendrogram of samples, ward.D2 method was used. Globin mRNA ratio of BC extraction_1, BC extraction_2 and BC extraction_3 was 9.76%, 19.46% and 23.23% respectively. (B) Coefficient of determination between two groups. (C) DEGs number in different comparation. Red box means up-regulated gene number, blue box means down-regulated gene number.

There were three gene expression patterns of DEGs (combine all DEGs in Figure 3C) as shown in Figure 4A. We observed 616 genes in Cluster 1 with the expression: RBC lysis > WB = BC extraction > PBMC isolation, and even some of the Cluster 1 genes were barely expressed in PBMC isolation. Cluster 2 had 413 genes and with the expression WB ≥ BC extraction > RBC lysis > PBMC isolation. Only 21 genes were observed in the Cluster 3 and expressed PBMC isolation > WB > BC extraction > RBC lysis, but all of these genes expressed much lower (Figure 4B). We expected pathway term enriched by DEGs would likely be due to differences in isolations, not biological variation. Genes in Cluster 1 were enriched in not only granulocyte-related functions, such as neutrophil activation and neutrophil degranulation, but also many immune activation pathways (Figure 4C). Cluster 2 genes were enriched in the functions related to red blood cells such as iron ion homeostasis and red blood cells differentiation (Figure 4C). Cluster 3 genes were too little to be enriched significantly in pathway terms. This results of enrichment of functional categories suggests that different treatment may be altering the gene expression profiles of the sample.

**Figure 4:**
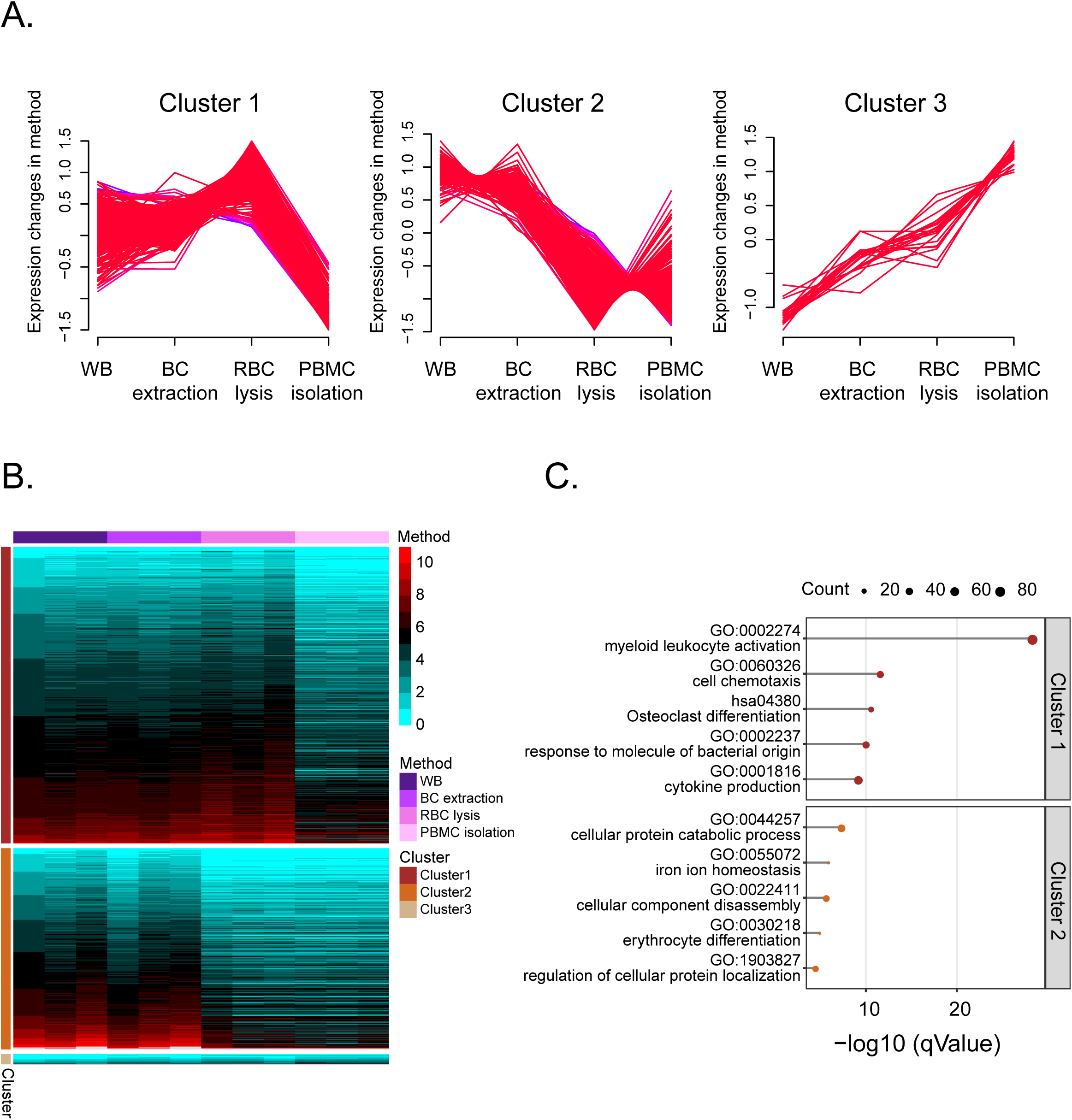
DEGs expression in WB and three leukocytes isolation methods. (A) MFUZZ used to cluster expression of DEGs (combined all DEGs in Figure 3C). Cluter1 has 616 genes, Cluster 2 has 418 genes. Cluster 3 has 21 genes. (B) Heatmap of DEGs’ expression in each sample. RPKM were transformed by log2(RPKM+1). (C) Gene annotation and analysis for DEGs in Cluster 1 and Cluster 2. Top 5 function clusters with their representative enriched terms were showed. Process enrichment analysis was carried out by Metascape (Methods).

## Discussion

*In vivo* changes in expression levels of leukocytes mRNAs can provide important information associated with different diseases or health status in large scale cohort study. There are always many samples to dispose at the same time in a large cohort study, so how to process sample efficiently, simply and stably is important. There is also a lack of clarity about which method to isolate leukocytes could reflect human physiological state preferably. This study provides an insight into the isolation effects on leukocytes gene expression profiles.

Firstly, our study found a significant influence of globin mRNA ratio on gene expression analysis among WB and three isolations, with a high globin mRNA ratio causing low transcriptome mapping rate and decrease the number of detected genes, which had similar results with early studies [22, 23]. RNA-seq analysis showed transcriptome of BC extraction was quite comparable to WB, further high globin mRNA and low gene detected ability were also observed in BC extraction, which had inconsistent aspiration of the BC extraction. The PBMC isolation contains monocytes and lymphocytes which are often used in immunology research [8] but showed relatively low expression of genes like FCGR3B and SLC44A2 which are markers of granulocytes (Figure S1). Compared three methods of isolate leukocytes from Table 1, we found RBC lysis removed much more globin mRNA than BC extraction, and remained more granulocyte information such as granulocytes than PBMC isolation. As globin mRNA will influence RNA-seq analysis and granulocyte play an important role in human immune system, RBC lysis was a much suitable method to isolate leukocytes and exclude globin mRNA to be able to perform full-scale gene expression profiling of leukocytes. Besides that, RBC lysis was much easier and less expensive to dispose than PBMC isolation, which had more advantage in large cohort studies. Research parameters and experimental conditions may limit the choice of methodology for blood processing and isolation before RNA-seq can be used to assess the transcriptome. Considering that study designs and objectives vary, further research might be needed for specific studies. Standard sample collection strategies need to be established before conducting research in order to minimize *ex vivo* change of blood and obtain meaningful and reproducible results.

**Table 1.** Summary of WB and three leukocytes isolation methods.

| Pretreatment |  | WB | BC | RBC | PBMC |
| --- | --- | --- | --- | --- | --- |
|  |  |  | extraction | lysis | isolation |
| **Experimental treatment** | Difficult of operation | + | ++ | ++ | ++++ |
|  | Operation time | + | ++ | +++ | ++++ |
|  | Reagent cost | + | + | ++ | ++++ |
|  | Equipment | No | Swinging<br>bucket | Swinging<br>bucket | Swinging<br>bucket |
|  |  |  | centrifuge | centrifuge | centrifuge |
|  | Globin mRNA ratio | ++++ | +++ | + | + |
| **Data analysis** | Mapping ratio | + | ++ | +++ | ++++ |
|  | Detected gene number | + | ++ | +++ | ++++ |
|  | Leukocytes related genes | +++ | +++ | +++ | + |
|  | Erythrocytes related genes | ++++ | ++++ | + | + |
| Legend: +low; ++moderate; +++high; ++++very high. |  |  |  |  |  |

## Research Funding

This project is supported by the National Natural Science Foundation of China (No.81300075), the Natural Science Foundation of Guangdong Province (No. 2014A030313795), the Shenzhen Municipal Government of China (No.JCYJ20170412152854656, JCYJ20180703093402288), the Shenzhen Peacock Plan (No. KQTD20150330171505310).

## Acknowledgments

We are grateful to the healthy volunteers who participated in the study. We are particularly grateful to professor Stephen Lye for valuable comments and professor Mark Johnson for proofreading. We also thank Dr Qing Zhou and Dr Zhongzhen Liu in our team give their comments on this study.

## Data accessibility

The data reported in this study are also available in the CNGB Nucleotide Sequence Archive (CNSA: https://db.cngb.org/cnsa; accession number CNP0000266).

## Author Contributions

Wen-Jing Wang and Hui Jiang designed research. Sujun Zhu and Juan Zeng collected samples. Yanru Xing, Xi Yang, Haixiao Chen and Yuan Chen processed samples. Yanru Xing, Xi Yang, Jinjin Xu and Fang Chen analyzed data. Yanru Xing, Wen-Jing Wang, Xi Yang and Hui Jiang wrote manuscript.

