## Supplemental Figure 1 for "Assessment of the gene expression profiles of leukocytes from different isolations"

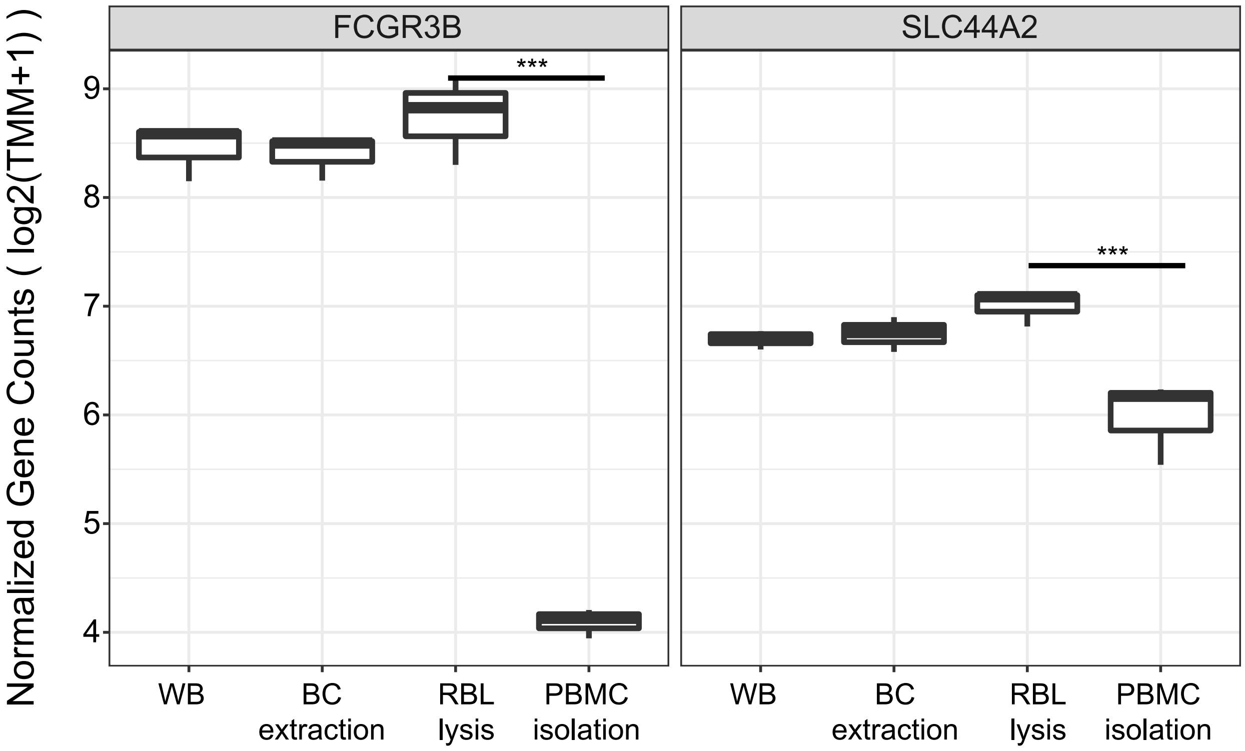


Fig. 7. Expression of FCGR3B and SLC44A2 in WB and three leukocytes isolation methods. FCGR3B and SLC44A2 which are markers of specific granulocytes cell types. *** p-adjust < 0.01.
